# NSMCE2, a Novel Super-Enhancer Regulated Gene, is Linked to Poor Prognosis and Therapy Resistance in Breast Cancer

**DOI:** 10.1101/2022.04.01.486781

**Authors:** Carolina Di Benedetto, Justin Oh, Zainab Choudhery, Weiquan Shi, Gilmer Valdes, Paola Betancur

## Abstract

In this study, we identified two novel super-enhancer associated genes: NSMCE2 and MAL2, highly upregulated in breast tumors, for which high RNA levels significantly and specifically correlate with breast cancer patients’ poor prognosis. To approach this, we took advantage of existing datasets containing super-enhancers associated genes identified in primary breast tumors and public databases comprising gene expression, genomic and clinical outcomes for patients diagnosed with breast cancer. Through in-vitro pharmacological super-enhancer disruption assays in breast cancer cells we confirmed that super-enhancers are involved in NSMCE2 and MAL2 transcript upregulation and through bioinformatics we found that high levels of NSMCE2 strongly associate with poor response to chemotherapy. This was observed especially for patients diagnosed with aggressive triple negative and HER2 positive tumor types. Finally, we showed that treating breast cancer cells with chemotherapeutic agents while simultaneously decreasing NSMCE2 gene expression by super-enhancer blockade or by directly silencing it, reduces cell viability thus increasing the effectiveness of chemotherapy. Our results indicate that moderating the transcript levels of the novel identified super-enhancer associated gene NSMCE2 could improve patients’ response to standard chemotherapy and, consequently, may improve disease outcome. In summary by mining existing public breast cancer datasets, our work demonstrates that searching for super-enhancer regulated genes and their association to patients’ survival and response to treatment, could be an effective method for identifying a signature of tumor specific -not frequently mutated, but super-enhancer dysregulated genes. Our approach offers a new avenue to identify novel biomarkers of poor prognosis and potential pharmacological targets for improving cancer treatment.

## Introduction

Breast cancer is the most diagnosed cancer type in women and the second leading cause of cancer-related female deaths in the US ^1^. Breast cancer is a heterogeneous disease but the identification of shared mechanisms that drive tumor progression has significantly improved patients’ treatment in the last decades. Breast tumors can be divided into subtypes using two parameters: (1) At the molecular level based on the protein expression of three receptors: estrogen receptor (ER), progesterone receptor (PR) and human epidermal growth factor receptor 2 (HER2) ^2^. Tumors that express hormone receptors ER and PR belong to the ER+PR+ subtype, while hormone receptor negative tumors with elevated HER2 levels belong to the HER2+ subtype. A third subtype called Triple Negative (TN) includes tumors that do not express ER, PR and HER2. (2) At the RNA level, based on gene expression profiles, breast tumors can be divided into 5 subtypes: Luminal A, Luminal B, HER2-enriched, Basal and Normal-like ^3–5^. The subtype classification of breast tumors using these parameters in combination with other clinicopathological characteristics such as age, tumor size, histological grade, and lymph node positivity, is clinically relevant for determining treatment recommendations ^6,7^. For instance, therapies aimed to block the activation of oncogenic receptors like ER and/or HER2 are available for the treatment of breast tumors expressing these receptors. While most patients with ER+ breast tumors receive endocrine therapy alone (e.g., tamoxifen), patients with HER2+ tumors are usually treated with anti-HER2 therapy plus chemotherapy. As these molecular targets are absent in TN tumors, TN breast cancer patients are frequently treated with chemotherapy.

Despite today’s current technology and advancements for the treatment of cancer, breast cancer related mortality, especially for TN breast cancer, remains high ^7,8^. One of the factors contributing to this is the lack of targeted therapies against TN tumors, which maintains TN cancers as an unmet clinical need ^9^. Another factor contributing to breast cancer recurrence and high mortality is tumor resistance to targeted therapies and/or chemotherapy, which favors the selection of less susceptible tumor cells over time ^10^. Thus, approaches aimed at identifying novel breast cancer targets of resistance, mostly for TN subtype, are urgently needed.

Super-enhancers (SEs) are long stretches of DNA containing clusters of enhancers that drive expression of genes of cell identity ^11,12^, and of other pivotal functions such as homeostasis ^13–15^. Because of several oncogenic pathways ^16,17^ and tumor immunosuppressive pathways ^18^ are also regulated by SEs, studying the role of SEs on the dysregulation of oncogenes has gained attraction over the last decade. SEs are typically enriched in acetylation at histone 3 lysine 27 (H3K27Ac) ^19^, a hallmark of open chromatin that allows DNA to be accessible to the transcriptional machinery. Lysine acetylation in histones is recognized by Bromodomain and Extraterminal (BET) proteins through epigenetic reader domains called bromodomains (BRDs). BRD4 is a member of the BET protein family that binds preferentially to hyperacetylated chromatin regions, hence SEs, to facilitate rapid gene transcription by linking enhancers or promoters to the TEFb (transcription elongation factor) complex ^20^. Previous studies have shown that disrupting BRD4 selectively affects the expression of genes that are associated with Ses ^16^. Since SEs are known to regulate the expression of several oncogenes, targeting these regulatory regions has been proposed as an anti-cancer therapy. This approach has the potential to simultaneously target multiple dysregulated pathways at the transcriptional level across cancer types.

In this work, by using publicly available databases containing gene expression data and clinical information obtained from cancer patients, we identified two SE-associated genes: NSMCE2 and MAL2 that are enriched in breast tumors when compared to non-cancerous tissue and that are linked to breast cancer patients’ poor prognosis. We show that pharmacological disruption of SE architecture using BET inhibitors decreases expression of these genes in vitro. Moreover, through analyses of tumor microarray data linked to therapy response of cancer patients, we found that high expression of NSMCE2 predicts poor response to chemotherapeutic drugs for patients diagnosed with breast cancer, especially patients diagnosed with aggressive TN or HER2+ tumors. Our results suggest that inhibiting NSMCE2 function by pharmacological inhibition or gene expression reduction (e.g., through SE blockade), could sensitize breast cancer cells to chemotherapy in specific cohorts of breast cancer patients that do not respond to standard antitumor drugs. This combinatorial approach has the potential to improve disease outcome for patients that do not survive after many years of cancer treatment and thus needs further investigation.

## Results

### Identification and characterization of candidate SE associated genes in breast cancer

In order to find genes associated to SEs, first we identified non-coding genomic regions carrying SEs from a dataset generated by applying the Ranking of Super-enhancers (ROSE) algorithm (https://bitbucket.org/young_computation/rose/) on H3K27Ac enrichment data. H3K27Ac data was obtained from chromatin immuno precipitation experiments followed by high throughput sequencing analyses (ChIP-Seq) performed in 4 breast cancer patient derived Xeno transplants (PDXs) from a previous study ^18^. We found that a tumor sample classified as hormone-receptor positive (ER+PR+) has 1194 DNA regions identified as SEs associated to 1128 genes, while the remaining three TN tumors contained 640, 527 and 651 SEs associated to 610, 458, and 587 genes, respectively (Fig 1A and Supplementary file). Our annotation analyses revealed that from the 3012 SEs identified, the majority are located either in chromosome 1 (14%), 8 (9%), 2 (9%), or 6 (9%) (Fig 1B, panel I). Since chromosome gene number widely varies across chromosomes, SE enrichment based on chromosome location was normalized by chromosome gene number (Fig 1B, panel II). Interestingly, when the number of genes per chromosome is considered, we found that SEs are also enriched in chromosome 8 (10%). From the total SE-associated genes, we identified 26 SE-associated genes common to all breast tumors analyzed (Fig 1A). Again, we found that most of these 26 SE associated genes are located in chromosomes 8 (27%) or 1 (23%) (Fig 1B, panel III). Moreover, when gene number per chromosome is considered, common breast cancer SE-associated genes remain particularly enriched in chromosome 8 (28%) (Fig 1B, panel IV). Given that chromosome 8 is known to contain many genes with tumor progression roles ^21,22^, our findings suggest SEs may be a frequent mechanism that dysregulates expression of oncogenes in breast tumors.

**Figure 1.**
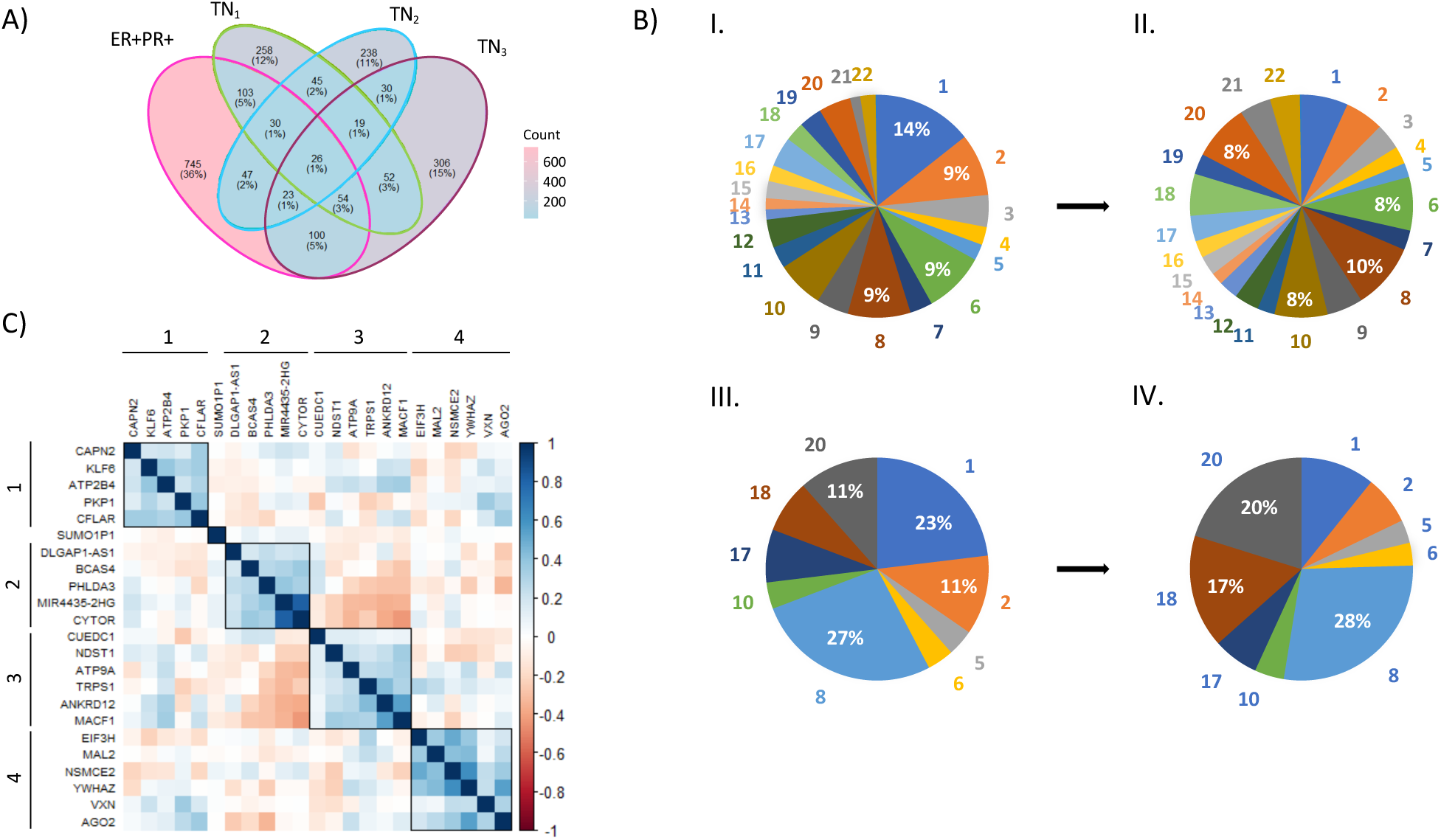
Identification and characterization of candidate SE associated genes in breast cancer. A) Venn diagram showing the distribution of DNA regions identified as SEs across the 4 breast PDX samples examined. 26 SE regions were found to be common to all samples. B) Pie charts for SEs enrichment based on chromosome location for the 4 breast PDX samples analyzed reveal that most SE regions (including those that are common to all the breast cancer samples analyzed) are enriched in chromosome 8 after normalization.B.I) Shows enrichment of all DNA regions identified as SEs. B.II) Shows enrichment of all DNA regions identified as SEs normalized by chromosome gene number. B.III) Shows enrichment of SE-associated genes common to all samples. B.IV) Shows enrichment of SE-associated genes common to all samples normalized by chromosome gene number. C) Heatmap showing gene expression correlations for the SE-associated genes found common to the 4 breast cancers analyzed, reveals 4 clusters of highly associated genes. This analysis was generated by using RNA-seq expression data from TCGA breast primary tumors (n = 1097). Chr = chromosome.

In order to investigate whether within these 26 SE-associated genes there are groups of genes that share mechanisms of gene regulation or interact in similar pathways, we performed a correlation matrix using hierarchical clustering of gene expression information from breast tumors. For this, we used bulk RNA-sequencing data (RNA-seq) from breast tumors available for 23 out of the 26 genes on The Cancer Genome Atlas (TCGA) database ^5^. Our correlation analysis identified four clusters of genes whose expression levels are significantly correlated (Figure 1C). From the hierarchical clustering we noted (1) a strong positive correlation between expression of CYTOR and MIR4435-2HG –two homolog non-coding RNA genes reported to be correlated with patients’ poor prognosis in glioblastoma and in low-grade glioma ^23^– and (2) an anticorrelation between expression of genes in the second and third clusters. Next, we performed a pathway enrichment analysis using Metascape ^24^, a web tool that combines functional enrichment, interactome (e.g., protein to protein, protein to DNA) analysis and gene annotation leveraging over 40 independent knowledgebases. Through Metascape we found that genes grouped in the third cluster are target genes for the transcription factor NF1, a tumor suppressor gene that has been reported to be associated with an increase in breast cancer risk when lost ^25^.

To further investigate the causes of gene expression correlation among the SE-associated genes in breast tumors, we analyzed co-occurrence of genetic alterations of genes within each cluster. For this, we used cBioPortal ^25^, an open-source platform that facilitates the interactive exploratory analysis and visualization of large-scale cancer genomics datasets. Using cBioPortal on TCGA genomic data we observed that highly co-expressed genes in cluster 1 (CAPN2, ATP2B4 and PKP1) and cluster 4 (EIF3H, MAL2, NSMCE2, YWHAZ, VXN and AGO2) have a frequency of genetic alterations between 8 to 10% and 7 to 15%, respectively, mainly due to co-occurring DNA amplifications (Supp Fig 1). CAPN2, ATP2B4 and PKP1 are located on chromosome 1 between the cytogenetic bands 1q32.1 and 1q41, while genes in cluster 4 (except for VXN) are located on chromosome 8 in a region spanning cytogenetic bands 8q22.3 to 8q24.3. Importantly, these regions on chromosomes 1 and 8 are frequently amplified in cancer disease ^5,26,27^. In line with our findings, recent reports show that SEs are frequently associated with genes located within large tandem duplications in breast cancers ^28^. Overall, our gene correlation analyses suggest that genes found in clusters 1 and 4 have SE-mediated enhanced transcription that help increase transcript levels of genes that are highly amplified and potentially involved in breast cancer development.

### High Gene Expression Levels of NSMCE2 and MAL2 SE-associated genes in breast tumors are linked to patients’ poor prognosis in breast cancer

Previously it has been reported that SEs can enhance the expression of tumor-associated genes. For instance, in T-cell acute lymphoblastic leukemia somatic mutations create a SE upstream of the TAL1 oncogene ^17^. Recently, by comparing breast tumor samples with healthy breast tissue samples we found that the immunosuppressive signal CD47 is transcriptionally upregulated by a SE ^18^. Thus, we decided to investigate whether the gene expression levels of our 26 SE-associated candidate genes are higher in breast tumors when compared to healthy samples by using the TCGA database and the Genotype-Tissue Expression (GTEX) study ^29^. TCGA compiles RNA-seq data of tumors or matched adjacent tissue (considered “healthy” tissue due to absence of cancerous characteristics) obtained from cancer patients, for whom clinical information is also available. Such information includes histological grade and survival data. GTEX provides open access to bulk RNA-seq gene expression data from non-diseased tissues obtained from donors. Mining these datasets we found RNA-seq data available for 21 out of the 26 SE-associated genes of our interest (RNA-seq data was not available for a pseudogene, and four non-coding RNA genes). From these 21 genes we found that 9 showed significantly higher expression in breast primary tumors when compared to matched adjacent normal tissue or to breast normal tissue samples from the GTEX study (Figure 2A and Supp Table 1). These data suggest that SEs could be driving the overexpression of these 9 genes in breast tumors.

**Figure 2.**
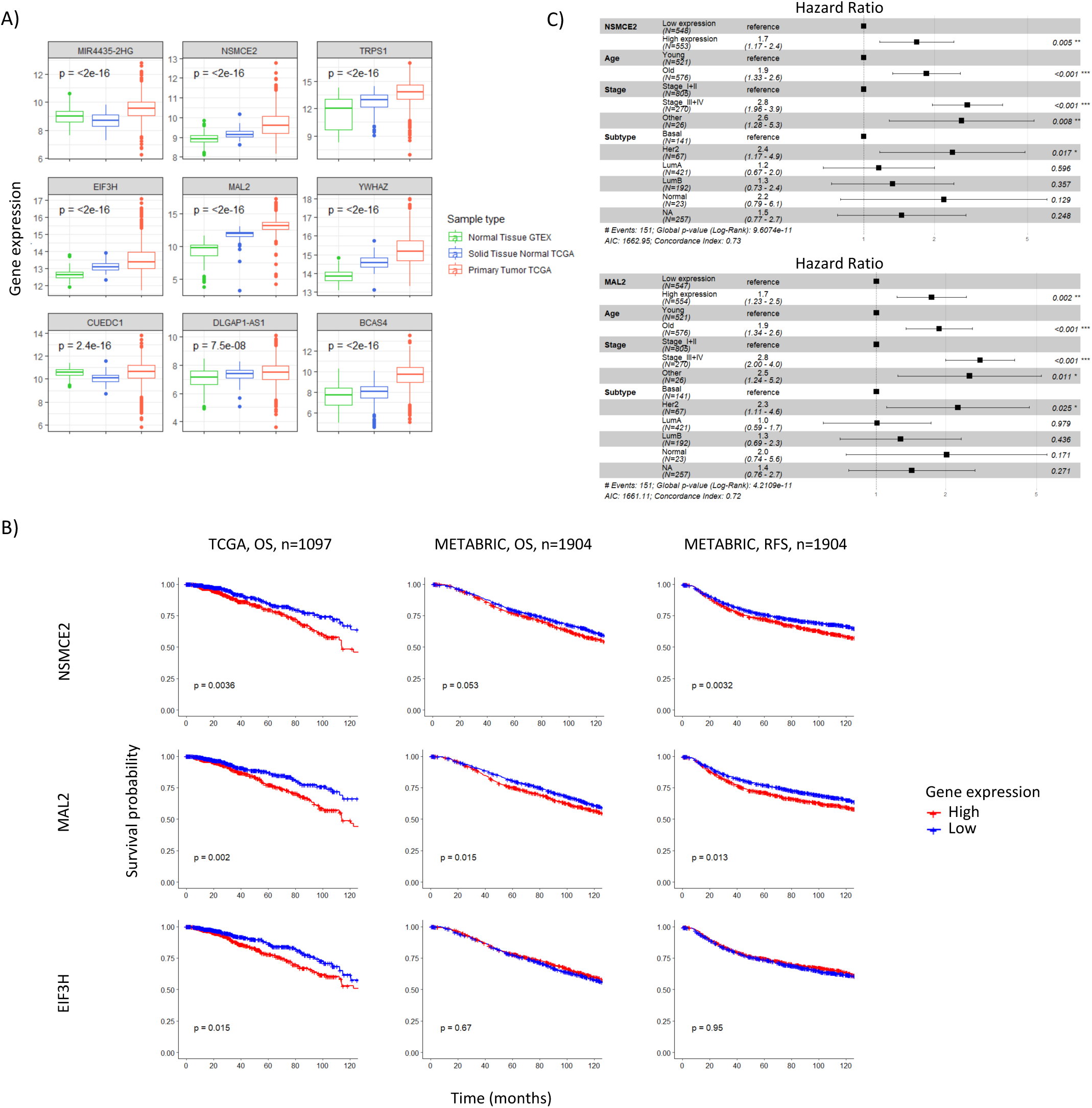
High NSMCE2 and MAL2 RNA expression in tumors is linked to poor prognosis of breast cancer patients. A) Box plots showing that gene expression levels are significantly higher in breast tumor samples when compared to normal samples for 9 SE-associated genes. Expression levels information for normal samples (Normal Tissue GTEX and Solid Tissue Normal TCGA) was obtained from GTEX and TCGA RNA-seq datasets, respectively. Expression levels from primary breast tumors (Primary Tumor TCGA) were obtained from TCGA RNA-seq datasets. Gene expression levels were compared by ANOVA. B) Univariate Kaplan-Meier plots showing breast cancer patients’ survival probability over time based on gene expression for each NSMCE2, MAL2, or EIF3H. These analyses were performed using breast primary tumors RNA-seq data mined from two independent datasets, TCGA (n = 1097) or METABRIC (n = 1904). Red and blue curves represent samples that show high and low gene expression levels, respectively, relative to the median expression value for each gene. High levels of either NSMCE2 or MAL2 were found to be significantly correlated with breast cancer patients’ poor prognosis in the two datasets. Plots were analyzed using the Log-rank test. OS = overall survival, RFS = relapse free survival. C) Multivariate Cox model of overall survival for NSMCE2 and MAL2 RNA expression in TCGA breast primary tumors including age, stage and tumor subtype as covariates show that high levels of either NSMCE2 or MAL2 are significantly correlated with patients’ poor prognosis.

Next, to investigate whether high levels of gene expression in breast tumors for any of the 26 SE-associated genes has clinical relevance, we performed a univariate Kaplan-Meir analysis using RNA-seq data linked to patients’ survival outcomes available for 1097 breast primary tumors through TCGA. We found that high transcript levels of 3 genes: NSMCE2, MAL2 and EIF3H, significantly correlate with worse overall survival in breast cancer patients (Log-rank test, p-value<0.05) (Fig 2B). To validate these results on an independent dataset, we performed the same univariate Kaplan-Meir analysis using the Molecular Taxonomy of Breast Cancer International Consortium (METABRIC) dataset, which contains microarray gene expression data from 2000 breast primary tumors along with long-term clinical follow-up ^30^. On the METABRIC dataset, high expression of MAL2 significantly associates with worse overall survival of breast cancer patients, while high NSMCE2 follows the same trend without reaching statistical significance (Fig 2B). However, on the METABRIC patient cohort high expression levels of both genes are significantly associated with reduced probability of relapse free survival –the length of time a cancer patient survives without any signs or symptoms of cancer after the end of primary treatment– thus, confirming that NSMCE2 and MAL2 expression are linked to poor prognosis in breast cancer. On the other hand, we did not observe a correlation between high EIF3H expression and breast cancer patients’ negative prognosis on the METABRIC dataset, even though both TCGA and METABRIC data sets do not differ on the probability of overall survival for their respective patients’ cohorts (Supp Fig 2A) and on other demographic characteristics ^31^. Since we found high levels of NSMCE2 and MAL2 gene expression highly correlate with patients’ negative prognosis, we focused our subsequent analyses on these two genes.

To further confirm that the association of high levels of gene expression with patients’ negative prognosis is independent of other risk factors (covariates), we constructed a multivariate Cox model of survival using TCGA data, which includes age, stage and RNA expression subtype as covariates. We observed that after adjusting for these clinical covariates, higher NSMCE2 and MAL2 gene expression levels remained significantly associated with patients’ poor prognosis (Figure 2C). NSMCE2 encodes a member of a family of E3 small ubiquitin-related modifier (SUMO) ligases, part of the structural maintenance of chromosomes 5/6 (SMC 5/6) complex. This protein complex plays key roles in DNA double-strand break repair by homologous recombination and in chromosome segregation during cell division ^32–34^. MAL2 is a multispan transmembrane protein required for transcytosis, an intracellular transport pathway used to deliver membrane-bound proteins and exogenous cargos from the basolateral to the apical surface ^35^.

Next, we investigated in other cancer types (apart from breast cancer) whether expression of NSMCE2 and MAL2 is higher in tumor samples when compared to normal tissue and whether high levels of any of these two genes associate with poor survival. To approach this, we compared RNA levels from an array of cancer samples from TCGA Pan-Cancer with RNA levels from corresponding normal tissue samples from GTEX and TCGA studies as we did for breast cancer. In line with what we observed in breast cancer, NSMCE2 or MAL2 are significantly higher in the cancer samples compared to the normal corresponding samples in most of the cancer types studied (Supp Fig 2B). When we asked whether their overexpression is also correlated with patients’ poor survival in other cancer types from the TCGA Pan-Cancer data set, we observed that high NSMCE2 RNA expression levels associates with worse overall survival in 6 other cancer types (apart from breast cancer). For MAL2, we found that while high gene expression is linked to shorter survival in 5 cancer types (including breast cancer), it is linked to better survival in 2 cancer types. Thus, although NSMCE2 and MAL2 are highly expressed in Pan-Cancer tumors, their association to patients’ negative survival outcomes is specific to certain cancer types, including breast cancer (Supp Table 2).

### NSMCE2 and MAL2 are regulated by SEs in ER+PR+, HER2+ and TN breast cancer cells

Since we found that SEs are associated to NSMCE2 and MAL2 expression in breast tumors, we hypothesized that in breast cancer cells, expression of these genes should be reduced when disrupting the SE enhancing function by blocking the binding of BRD4 to SEs with the inhibitors JQ1 or IBET151 ^36,37^. To test this, we used 5 breast cancer cell lines representing different breast cancer subtypes: MCF7 (ER+PR+), HCC1954 (HER2+), and BT549, MDAMB231 and Hs578T (TN). We also chose these cell lines because NSMCE2 and MAL2 are amplified in all except in MDAMB231 (visualized using cBioPortal on data from the Cell Line Encyclopedia ^38^, Supp Fig 3A) which recapitulates our observation in breast cancer patients’ data sets where these two genes were frequently amplified.

Treatment of ER+PR+, HER2+, and TN cancer cell lines with non-toxic concentration of either JQ1 or IBET151 (Supp Fig 3B) had the following effects on gene expression levels (Figure 3): We observed a reduction on NSMCE2 RNA in all cell lines treated with BET inhibitors, confirming regulation of NSMCE2 by SEs. MAL2 expression was only reduced by BET inhibition in cell lines representing HER2+ and TN breast cancer subtypes, but it was curiously increased after 24h treatment in the TN BT549, perhaps as an indirect effect of BET inhibition on downregulating MAL2 repressors. In summary, pharmacological blockade of SEs confirmed NSMCE2 and MAL2 are associated with SEs in breast cancer cells.

**Figure 3.**
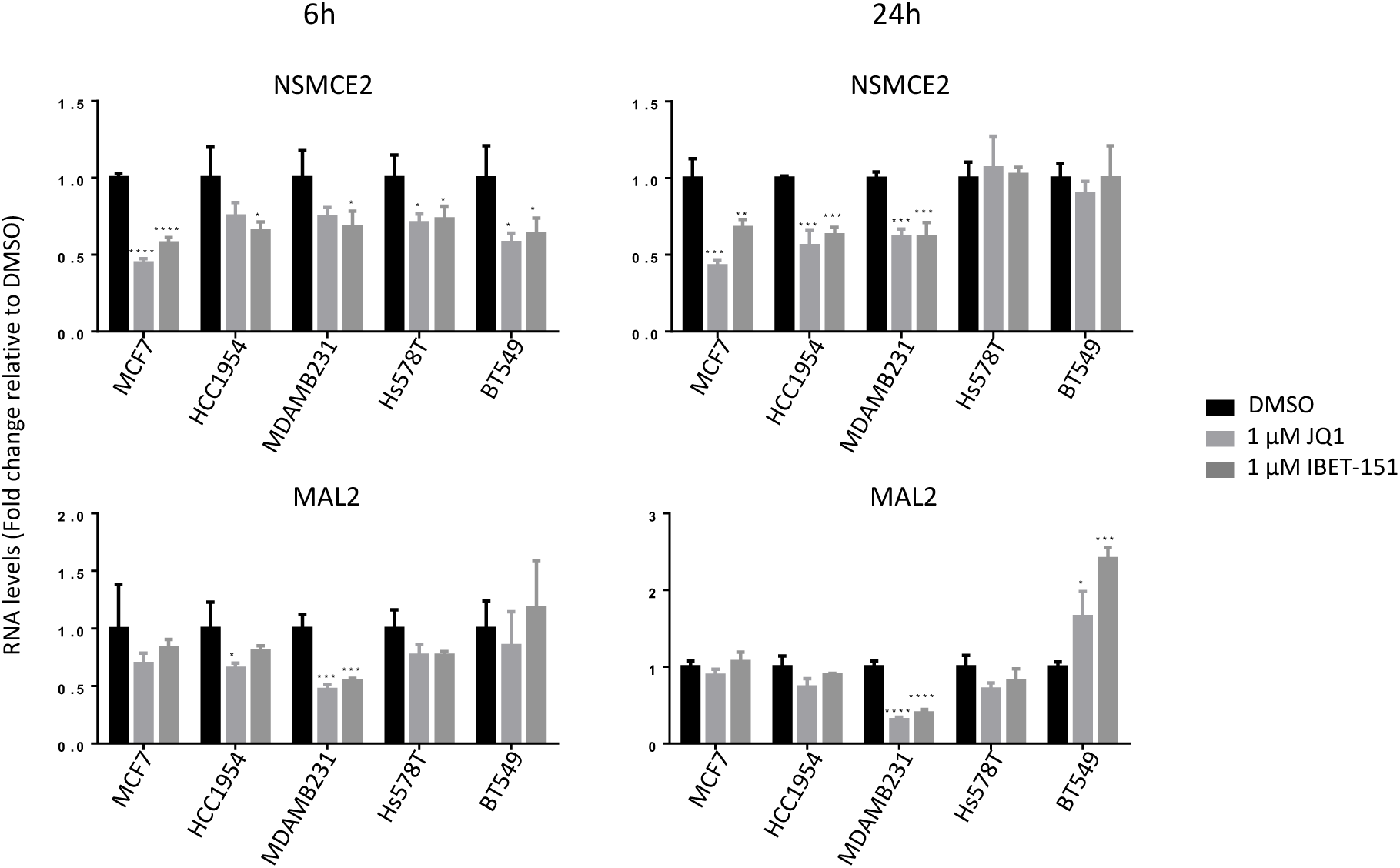
NSMCE2 and MAL2 RNA expression is regulated by SEs in breast cancer cells. Blocking BRD4 binding to SEs with BET inhibitors reduces gene expression for NSMCE2 and MAL2 starting at 6 hours in most breast cancer cell lines tested. MCF7, HCC1954, MDA-MB-231, Hs578T and BT-549 breast cancer cells were treated with vehicle (DMSO), 1 μM JQ1 or 1 μM IBET-151 for 6 and 24 hours. After treatment, changes in gene expression levels were analyzed by qPCR. ANOVA followed by Dunnett’s multiple comparison test, *P < 0.05; **P < 0.01; ***P < 0.001; ****P < 0.0001 was performed.

### High Gene Expression Levels of NSMCE2 correlate with breast cancer patients’ poor response to chemotherapy

Given that our data shows that high NSMCE2 and MAL2 expression are associated with breast cancer patients’ poor survival outcomes, we next investigated if high levels of these markers could also be negatively associated with response to standard cancer therapies. To investigate this, we used the online platform ROC Plotter (http://www.rocplot.org), which integrates multiple transcriptome-level gene expression datasets available through GEO into a single database containing information for 3104 breast cancer patients linked to corresponding response data for a range of treatments, including endocrine therapies, anti-HER2 therapies and chemotherapeutic drugs ^39^. The transcriptome data is obtained from patient’s biopsies prior to treatment and patients are grouped into responders and non-responders for a given treatment based on clinical characteristics. Responder and non-responder patients are compared via two diverse statistical approaches: The Mann–Whitney test (non-parametric T-test) and the Receiver Operating Characteristic (ROC) test. The ROC test measures how much ‘the gene expression level’ model is capable of distinguishing between the two classes: responders and non-responders. ROC plots are useful to compute the Area Under the Curve value (AUC), which shows the predictive power of the gene. The higher the AUC value, the better the model is at distinguishing between patients who are responders versus non-responders. For cancer biomarkers with potential clinical utility, the AUC value should be higher than 0.6, while AUC values higher than 0.7 are obtained by top quality cancer biomarkers ^39^.

We focused our analysis on breast cancer patients who received chemotherapy due to the larger sample size in the dataset compared to that for patients who received endocrine or anti-HER2 therapies (n=507 versus n=160 and n=151, respectively). Using ROC Plotter, we found that high levels of NSMCE2 gene expression correlate with lower pathological complete response to chemotherapy of breast cancer patients (Fig 4A, Mann-Whitney test p-value=0.000032 and AUC=0.617). Moreover, we found that high levels of NSMCE2 gene expression strongly correlate with poor response to chemotherapy for patients diagnosed with grade III breast cancer (Fig 4B, Mann-Whitney test p-value=0.00099 and AUC=0.655). We also performed the same analyses for the 21 SE-associated genes we found in breast cancer and for which gene expression data is available on ROC plotter. We did not see significant correlation (AUC<0.6, Mann-Whitney test p-value<0.05) between high levels of gene expression and breast cancer patients’ poor response to chemotherapy for any of the other SE-associated genes analyzed (including MAL2), when analyzing the entire database or when focusing on Grade III breast cancer (Supp Table 3). This indicates that the observed correlation between high NSMCE2 gene expression levels and poor response to chemotherapy is specific.

**Figure 4.**
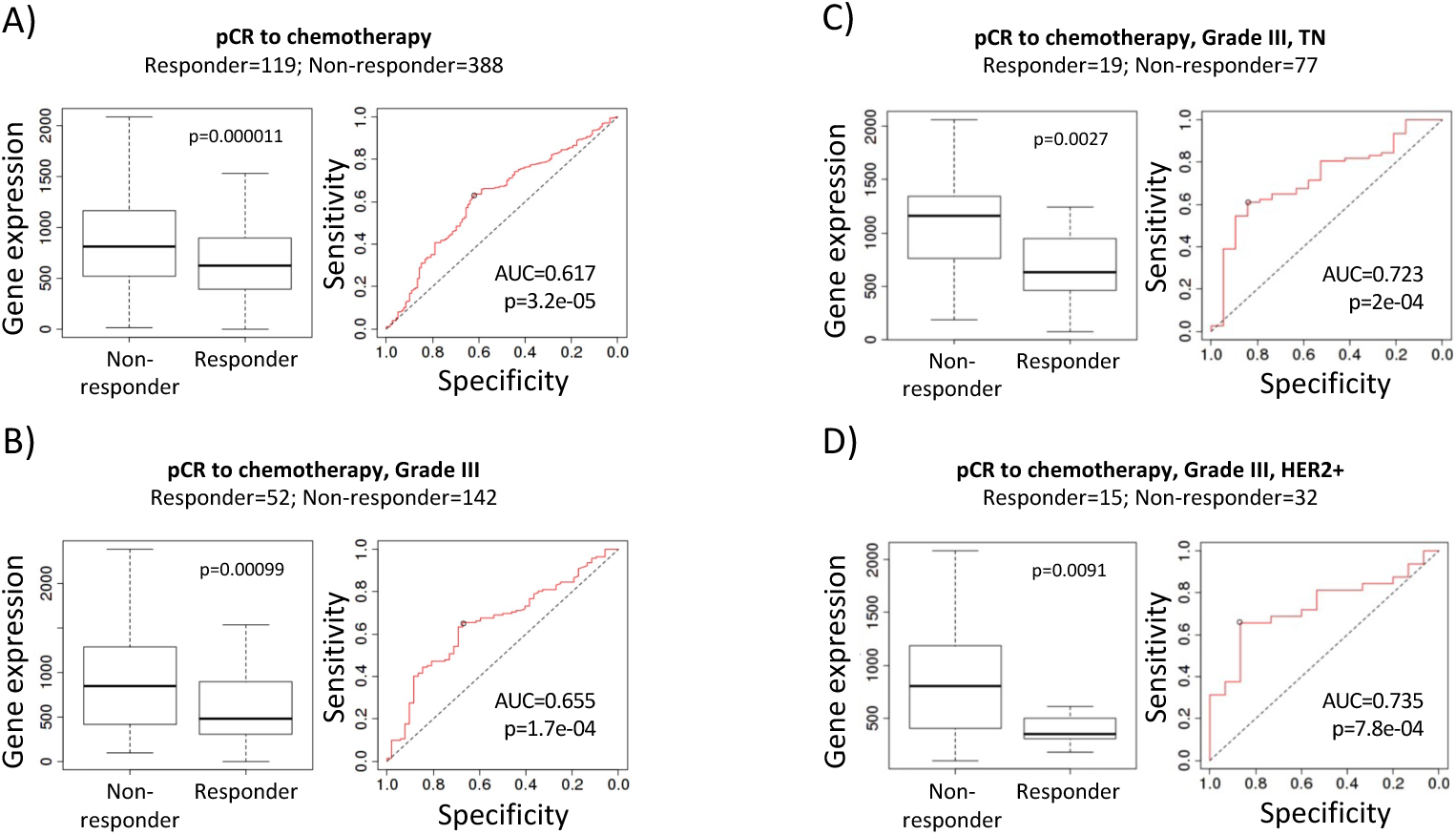
High NSMCE2 expression correlates to breast cancer patients’ poor response to chemotherapy. Left panels are box plots showing NSMCE2 gene expression levels in non-responder versus responder patients. Right panels are receiver operating characteristic (ROC) plots showing the specificity and sensitivity of NSMCE2 gene expression as a predictor for response to chemotherapy in: A) breast cancer (n = 507), B) Grade III breast cancer (n = 194), C) Grade III TN breast cancer (n = 96), and D) Grade III HER2+ breast cancer (n = 47). In the four cases, non-responder patients have significantly higher NSMCE2 expression levels compared to the responder group and the AUC values indicate that NSMCE2 gene expression level is capable of distinguishing between responder and non-responder patients. Analyses shown in this figure were performed using ROC Plotter, Mann-Whitney U test and Receiver Operating Characteristic test. pCR = pathological complete response.

Since the drug regimen for breast cancer patients is highly determined by the tumor molecular subtype, we next analyzed the correlation between high NSMCE2 gene expression levels with response to chemotherapy for each ER+PR+, HER2+ or TN tumor subtypes. Here, we observed a stronger correlation between high levels of NSMCE2 and patients’ poor response to chemotherapy when the tumors were classified as grade III TN or grade III HER2+ tumors (Fig 4C and D, AUC= 0.723, Mann-Whitney test p-value=0.0027 and AUC= 0.735, Mann-Whitney test p-value=0.0091, respectively). Thus, our results indicate that breast cancer patients with high NSMCE2 RNA expression in aggressive breast tumors of either TN or HER2+ molecular subtype, respond poorly to chemotherapeutic drugs. Since the AUC values obtained for NSMCE2 are usually values observed for top quality cancer biomarkers ^39^, our findings suggest that NSMCE2 expression could potentially be used to pinpoint at a group of patients (especially those diagnosed with grade III TN or HER2+ breast cancer) that may not respond to chemotherapy.

Lastly, using the ROC Plotter tool, we found that NSMCE2 high expression levels do not correlate with patients’ response to therapy for ovarian cancer, colorectal cancer, and glioblastoma ^40,41^ (data not shown), thus, confirming that high levels of NSMCE2 gene expression are specifically associated with breast cancer patients’ poor response to chemotherapy.

### Lowering NSMCE2 transcript levels sensitizes breast cancer cells to chemotherapeutic agents

So far, our work shows that (1) NSMCE2 and MAL2 genes are regulated by SEs at the transcript level in breast cancers; (2) the expression of these genes can be reduced by BET inhibition; and (3) high NSMCE2 expression correlates to poor response to chemotherapy in grade III TN or HER2+ breast cancer patients. The latter result indicates that our findings may have important clinical implications. Most chemotherapeutic agents work by activating the program of apoptosis in cancer cells through DNA damage induction or through cell cycle inhibition. Since NSMCE2 is known to be required for DNA damage repair at different steps of the process and for chromosomal segregation during mitosis ^32–34,42–44^, NSMCE2 may inhibit chemotherapy-induced apoptosis, thus contributing to the therapy resistance we observed in breast cancer patients showing high NSMCE2 gene expression. To test whether reduction of NSMCE2 transcript levels by SE disruption can sensitize breast cancer cells to chemotherapy-induced apoptosis, we treated breast cancer cells with BET inhibitor JQ1 and the standard chemotherapeutical drugs doxorubicin and paclitaxel –two antitumor agents widely used for the treatment of breast cancer–. Doxorubicin is part of the anthracycline group of chemotherapeutic agents that exert antitumor action by both DNA intercalation and inhibition of the enzyme topoisomerase II, which results in DNA damage during DNA replication, and the eventual induction of apoptosis. Paclitaxel, on the other hand, is a class of taxanes, an antimitotic drug that affects the stabilization of microtubules, thus, disrupting the cell ability to divide ^45^.

Even though we observed NSMCE2 expression is reduced by SE blockade in all the breast cancer cell lines tested, we decided to use TN breast cancer cells, MDAMB231, BT549 and Hs578T, and the HER2+ cells HCC1954 for our experiments and analyses, since we found that high NSMCE2 gene expression levels significantly associate with poor response to chemotherapy for patients diagnosed with TN or HER2+ breast cancer. Initially, cells were pre-treated with the BET inhibitor JQ1 for 3 hours to block SEs transcriptional function (this is a window of time we have previously found successfully inhibits SE function). After the 3 hours, cells were treated with JQ1 in combination with several concentrations of either of the chemotherapeutic drugs. We assessed cell viability by MTT assays. For all cell lines tested we observed that the reduced cell viability resulting from the combined treatment is highly dependent on the chemotherapeutic agent used and on its concentration (Supp Fig 4A). For instance, combination of JQ1 and doxorubicin has an additive effect on cell survival reduction when using doxorubicin at 1 µM. Such additive effect is gradually lost at higher concentrations of doxorubicin, where we see strong decline in cell viability regardless of SE inhibition by JQ1, probably due to high toxicity and massive DNA damage caused by doxorubicin. On the other hand, for any of the breast cancer cell lines tested, paclitaxel effectiveness at reducing the number of viable cells did not improve when given in combination with JQ1, in fact, an antagonistic effect was observed upon combination of these drugs. Next, to replicate the scenario we observe when analyzing the correlation of NSMCE2 gene expression levels with response to chemotherapy in breast cancer patients (where we see low NSMCE2 expression levels correlate with good response to chemotherapy), we pre-treated cells with JQ1 for 24 hours in order to allow enough time to reduce expression levels of SE-associated genes (in particular for NSMCE2, Fig 3) before doxorubicin treatment. Here, we also observed an additive effect on cell viability reduction in response to doxorubicin when NSMCE2 levels are lowered by JQ1 treatment (Supp Fig 4B). Since treating cells with JQ1 prior to doxorubicin treatment increases doxorubicin effectiveness at reducing cell viability, we wonder whether this reduction in viable cell numbers is due to increased apoptosis. To assess this, we measured apoptosis by Annexin V staining using flow cytometry. Breast cancer cells were pretreated with JQ1 for 24 hours prior to the addition of JQ1 and/or chemotherapeutic drugs. As shown in figure 5A, doxorubicin highly increases apoptosis of BT549 cells and of Hs578T to a lower extent. Moreover, the combination of doxorubicin with JQ1 treatment strongly synergizes to induce apoptosis in both breast cancer cell lines. On the contrary, neither paclitaxel alone nor in combination with JQ1 increase apoptosis of the treated breast cancer cell lines (Figure 5B).

**Figure 5.**
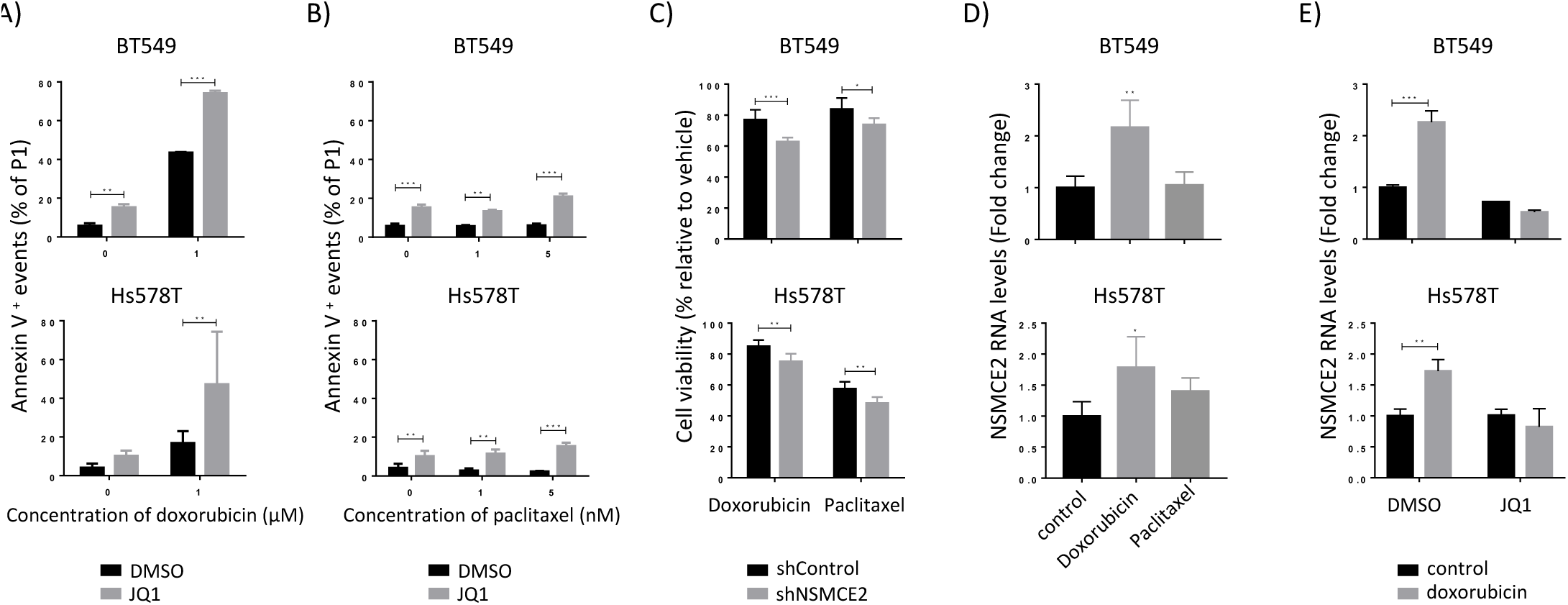
Lowering NSMCE2 transcript levels sensitizes breast cancer cells to chemotherapeutic agents. A) Measuring the effect of SE blockade and doxorubicin treatment on apoptosis of breast cancer cell lines by Flow Cytometry shows that the combined treatment strongly synergizes to induce apoptosis in both breast cancer cell lines BT549 and Hs578T were preincubated with vehicle (DMSO) or 1 μM JQ1 for 24 hours, after which culture medium was replaced by fresh medium containing vehicle or 1 μM JQ1 +/- 1 μM doxorubicin. After 48 hours, apoptotic cells were quantified by Annexin V staining. Two Way ANOVA followed by Bonferroni’s multiple comparison test was performed, **P<0.01, ***P<0.001. B) Measuring the effect of SE blockade and paclitaxel treatment on apoptosis of breast cancer cell lines by Flow Cytometry shows that neither paclitaxel alone nor in combination with JQ1 increase apoptosis of the treated breast cancer cell lines. BT549 and Hs578T were preincubated with vehicle (DMSO) or 1 μM JQ1 for 24 hours, after which culture medium was replaced by fresh medium containing vehicle or 1 μM JQ1 +/- 5 nM paclitaxel. After 48 hours, apoptotic cells were quantified by AnnexinV staining. Two Way ANOVA followed by Bonferroni’s multiple comparison test was performed, **P<0.01, ***P<0.001. C) Knocking down NSMCE2 by shRNAs in breast cancer cells shows that reducing NSMCE2 levels sensitizes cells to chemotherapeutic drugs, improving their effect on cell viability reduction. BT-549 and Hs578T transduced with shControl or shNSMCE2 were incubated with 0.5 μM doxorubicin for 48 hours or 5 nM paclitaxel for 72 hours. After treatment, viable cells were quantified by the MTT assay. Plots show the percentage of cell viability for the treatment relative to the vehicle control condition. Two Way ANOVA followed by Bonferroni’s multiple comparison test was performed, *P<0.05, **P<0.01, ***P<0.001. D) NSMCE2 RNA expression levels measured by qPCR upon chemotherapeutic drug treatment of breast cancer cell lines, show that NSMCE2 transcript expression increases upon doxorubicin treatment but not upon paclitaxel in BT549 and Hs578T treated cancer lines. Cells were treated with 1 μM doxorubicin or 5 nM paclitaxel for 24 hours. ANOVA followed by Dunnett’s multiple comparison test was performed, *P<0.05, **P<0.01. E) Analysis of NSMCE2 RNA expression upon SE blockade and doxorubicin treatment demonstrates that the NSMCE2 transcript increase observed after doxorubicin treatment is mediated by SEs in BT549 and Hs578T. BT549 and Hs578T were treated with 1 μM JQ1 and/or 1 μM doxorubicin for 24 hours and NSMCE2 RNA expression was determined by qPCR. Two Way ANOVA followed by Bonferroni’s multiple comparison test was performed, **P<0.01.

In summary, our results demonstrate that JQ1 synergizes with doxorubicin to induce apoptosis suggesting that reducing NSMCE2 expression by SE inhibition may contribute to the increase in apoptosis of the breast cancer cell lines tested. To confirm that NSMCE2 downregulation by JQ1 SE blockade is specifically contributing to the stronger effect observed on cell viability reduction, we next generated BT549 or Hs578T cells stably expressing a short hairpin RNA against NSMCE2 (shNSMCE2) or control (shControl) to knockdown gene expression. After checking NSMCE2 silencing was efficient (Supp Fig 4C), we performed MTT assays to measure cell viability upon chemotherapeutic drug treatments. Here, we found that silencing NSMCE2 in both cell lines significantly contributed to a greater reduction in cell viability after 48 hours of incubation with doxorubicin. Knocking down NSMCE2 had a significant but smaller contribution to cell viability reduction after 72 hours of paclitaxel treatment (Figure 5C). Taken together, these results show that reducing NSMCE2 levels sensitizes breast cancer cells to chemotherapy, thus implying that NSMCE2 high levels contribute resistance to chemotherapeutic agents.

Since NSMCE2 is required for DNA damage repair, we hypothesized that more NSMCE2 is produced at the transcriptional level upon chemotherapy-induced DNA damage. Indeed, we observed that NSMCE2 RNA levels significantly increase 24 hours after doxorubicin treatment in BT549 and Hs578T, but not upon paclitaxel treatment (Figure 5D). Given that NSMCE2 is associated with SEs in both cell lines, we next investigated if doxorubicin-induced NSMCE2 upregulation is mediated by SEs. Treatment of breast cancer cells with SE inhibitor JQ1 abrogates the effect of doxorubicin in NSMCE2 upregulation, indicating that NSMCE2 upregulation by doxorubicin is mediated by SEs (Figure 5E). Collectively, our results show that NSMCE2 expression is transcriptionally upregulated upon doxorubicin treatment through SEs, and this could be a factor contributing resistance to chemotherapy during breast cancer treatment. In sum, our experiments show that the correlation between high NSMCE2 expression levels in breast tumors and cancer patients’ poor response to chemotherapy is due in part to increased resistance to chemotherapeutic drugs driven by NSMCE2.

## Discussion

In this work, we mined publicly available cancer data sets containing cancer patients genomic and clinical information and performed experimental perturbations to identify novel genes erroneously upregulated in breast cancer by SEs and that are associated to patients’ poor outcome and resistance to cancer therapies. By using this approach, we identified two highly upregulated genes in tumors, NSMCE2 and MAL2, for which high levels of gene expression significantly correlate with breast cancer patients’ negative prognosis. Through pharmacological SE disruption assays performed in breast cancer cell lines we experimentally confirmed that SEs regulate the expression of NSMCE2 or MAL2. Importantly, by mining a database containing breast cancer treatment and response information, we found that high levels of NSMCE2, specially in grade III TN and HER2+ breast cancers, are strongly correlated with patients’ poor response to chemotherapy. Accordingly, we experimentally showed that reducing NSMCE2 gene expression levels increases the effectiveness of chemotherapeutic agents.

Previously it was reported that NSMCE2 is involved in maintaining telomeres length in cancer cell lines, in this manner preventing telomere-mediated cell cycle arrest and the activation of the senescence pathway ^46^. In a breast cancer cell line, NSMCE2 depletion has been reported to prevent cell cycle progression ^47^. Similarly to NSMCE2, MAL2 has a role in cancer disease, but through modulation of the immune system. MAL2 has been previously shown to reduce tumor cells’ antigen presentation by promoting the endocytosis of tumor antigens ^48^. Moreover, in breast cancer Jeong and colleagues showed that MAL2 leads to HER2 accumulation on the cell surface and enhanced HER2 signaling ^49^. These known tumor promoting roles for NSMCE2 and MAL2 suggest that SE-driven overexpression of these two genes can support a variety of tumor promoting processes in breast cancers cells. Since our results demonstrate that NSMCE2 and MAL2 gene expression levels are reduced by pharmacological SE-blockade, targeting SEs could work as a therapeutic approach to reduce these tumor promoting processes.

Through mining genomic datasets, we also found that NSMCE2 and MAL2 are frequently amplified in breast cancer patients and in breast cancer cell lines. Recently it was reported that SEs are frequently found on amplified DNA regions in tumor cells ^28^. DNA amplification is known to contribute to cancer progression by mediating overexpression of oncogenes or genes that promote cancer therapy resistance ^50^. Thus, frequent copy number gains of the NSMCE2 and MAL2 loci in breast cancer suggest that higher gene expression levels for these genes indeed contribute to tumor progression. Since we also demonstrated that downregulation of NSMCE2 and MAL2 gene expression can be achieved by blocking SEs, collectively these data suggest that both mechanisms (i.e., SE-driven dysregulation and gene amplification) can increase expression of these oncogenes in tumor cells. Thus, reducing the expression of these genes through SEs perturbation could have potential therapeutic value.

Currently, several ongoing clinical trials are studying SE disruption through BET inhibition, alone or in combination with other therapies, for the treatment of both hematological malignancies and solid tumors ^51^. However, SEs are also involved in the control of processes in healthy cells, bringing concerns about toxicity when using SEs inhibitors as treatment. Some common adverse effects observed in current clinical trials include thrombocytopenia (most frequent dose-limiting toxicity), nausea, diarrhea, and fatigue ^52–54^. For this reason, synergistic therapies have gained interest because they lead to increased efficacy with lower drug dose, thus reducing toxicities associated with higher concentrations. Following this approach BET inhibitors have been shown to sensitize tumor cells to antitumor drugs when tested preclinically. For instance, it has been shown that JQ1 synergizes with daunorubicin, an anthracycline chemotherapeutic drug, to induce apoptosis in acute myeloid leukemia cells ^55^. More recently, a combination of JQ1 and the CDK4/6 inhibitor palbociclib has been found to synergistically inhibit cell growth through induction of cell cycle arrest in TN breast cancer cell lines ^56^.

Our results show that JQ1 sensitizes breast cancer cells to the chemotherapeutic drug doxorubicin, by increasing apoptosis and by exerting an additive effect on cell growth inhibition when given in combination. Although this additive effect on cell viability reduction has been reported for JQ1 and doxorubicin treated primary osteosarcoma cells ^57^, we are the first to report this effect on breast cancer. On the contrary, we did not see an increase on apoptosis when breast cancer cells were treated by combining JQ1 with paclitaxel. We speculate that the increase in apoptosis observed upon JQ1 and doxorubicin combination depends mainly on the mechanism of action of the chemotherapeutic agent. For instance, while doxorubicin induces DNA damage, JQ1 through SE inhibition may be downregulating the expression of antiapoptotic genes or genes involved in DNA damage repair (including NSMCE2), thus leaving the cells with no other choice but to activate the apoptotic pathway upon doxorubicin treatment. In line with this, we found that NSMCE2 RNA expression increases upon doxorubicin treatment in breast cancer cells, suggesting that NSMCE2 upregulation could be required to overcome doxorubicin induced DNA damage. This upregulation is inhibited by BET inhibitors and, in consequence, mediated by SEs. Importantly, by generating NSMCE2 knockdown breast cancer cell lines, we showed that NSMCE2 plays a key role in resistance to doxorubicin treatment, probably due to its DNA repairing function, which reduces doxorubicin’s DNA damaging effect and prevents the activation of the apoptotic program. In line with our results, depletion of the NSMCE2 homologous in yeast, MMS21, was reported to hypersensitize cells to doxorubicin induced apoptosis ^58^. Altogether, our experimental findings imply that combining SE blockade with doxorubicin can be an effective therapy to treat breast tumors that express high levels of NSMCE2. Importantly, our computational analyses show that high NSMCE2 gene expression levels correlate to patients’ poor response to chemotherapy, especially in grade III TN and HER2+ breast cancers. Although we ignore the epigenetic modifications as well as the gene expression changes occurring in response to treatment in these breast tumors, this correlation also points at the role of NSMCE2 on resistance to chemotherapy. Taken together, our results suggest that reducing NSMCE2 levels could help to overcome resistance to DNA-damage inducing therapies in specific cohorts of patients.

In conclusion, our work demonstrates for the first time that in breast tumors NSMCE2 and MAL2 are dysregulated by SE, frequently amplified and overexpressed, and that their expression levels are associated to patients’ poor survival. We propose that reducing gene expression of these genes by SE inhibition or directly inhibiting their activities through pharmacological inhibition, could be effective strategies to prevent their tumor progression activities and thus needs to be further explored.

## Materials and Methods

### Analyses of cancer patients’ datasets

TCGA breast cancer dataset was downloaded from the UCSC XenaBrowser Platform ^59^. METABRIC data was downloaded from cBioPortal ^25^. UCSC Toil RNAseq study containing TCGA and GTEX gene expression was used to compare gene expression in tumor and normal tissues ^29^. Gene expression correlation matrix for SE-associated genes was done by hierarchical clustering using the package ‘corrplot’ on RStudio. Genetic alterations in breast cancer patients on the TCGA dataset or in breast cancer cell lines on The Cell Line Encyclopedia were analyzed using cBioPortal. Kaplan-Meier and Cox analyses were performed using the packages ‘survival’ and ‘survminer’ on RStudio and samples were divided in two groups based on the median gene expression. Kaplan-Meier curves were compared using the log-rank test. Association of gene expression and response to therapy in breast cancer patients was done using ROC Plotter ^39^.

### Cell culture

MCF7, HCC1954, MDAMB231, BT549 and 293T cell lines were obtained from ATCC. Hs578T were kindly provided by Dr. Mary Helen Barcellos-Hoff at UCSF. MCF7, HCC1954, BT549 and Hs578T cells were cultured in RPMI medium (Gibco), while MDAMB231 and 293T cells were cultured in DMEM medium (Gibco). Media were supplemented with 10% heat inactivated FBS (Atlanta Biologicals), 100 units/mL Penicillin and 100 µg/mL Streptomycin (Gibco). BT549 and Hs578T were also supplemented with 0.01 mg/mL insulin (Sigma). Cells were grown at 37°C in a humified atmosphere at 5% CO_2_.

### Drugs

For BET inhibition, cells were treated with JQ1 (Sigma) or IBET-151 (GSK1210151A, Selleckchem) dissolved in DMSO. For treatment with chemotherapeutic drugs, Doxorubicin hydrochloride (Sigma) dissolved in water and Paclitaxel from Taxus brevifolia (Sigma) dissolved in DMSO were used at the indicated concentrations. Diluents were used as vehicle controls.

### MTT Assay

Cells were seeded in 96-well plates (at 7000 cells/well) and cultured overnight. Then, cells were exposed to drug treatments for the indicated times. MTT (Sigma) was added to each well following manufacturer’s instruction. After incubation for 4 hours at 37°C, formazan crystals were dissolved in DMSO and absorbance was read at 570 nm on a Biotek plate reader. Combination Indexes (CI) were calculated for JQ1 in combination with each chemotherapeutic drug using the Response Additivity approach ^60^ as follows:

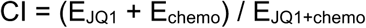

Where E_JQ1_ is the effect of JQ1 on cell viability reduction, E_chemo_ is the effect of a particular concentration of chemotherapeutic drug on cell viability reduction, and E_JQ1+chemo_ is the effect of the combined treatment on cell viability reduction for that concentration of chemotherapeutic drug. CI = 1 indicates additive effect, CI < 1 indicates synergistic effect, and CI > 1 indicates antagonism.

### RNA extraction, cDNA synthesis and qPCR

Total RNA was extracted using RNeasy Plus kits (Qiagen). cDNA was reversed transcribed using SuperScript III First-Strand Synthesis SuperMix (Invitrogen) and then amplified on the QuantStudio 5 Real Time PCR System (Applied Biosystems). Specific primers designed to amplify the desired gene were combined with cDNA and Platinum® SYBR® Green qPCR SuperMix-UDG (Invitrogen) following the manufacturer’s instructions. Primers used in this study are:

NSMCE2: AGGACGCCATTGTTCGCAT, GCTACAGCCAATTTGAGGGCA

MAL2: TCCGTGACAGCGTTTTTCTTTTC, TGCTTCCAATAAAAAGGCTCCAA

β-Actin: TCCCTGGAGAAGAGCTACG, GTAGTTTCGTGGATGCCACA

### Apoptosis assay by Flow Cytometry

Cells were collected, washed with HBSS and stained with APC Annexin V (Biolegend, Catalogue number 640919) in Annexin V binding buffer (Biolegend) for 30 minutes at room temperature. After washing, AnnexinV-positive events were quantified using a Northern Light cytometer (Cytek). Data were analyzed on SpectroFlo software (Cytek).

### Lentiviral transduction for NSMCE2 knockdown

NSMCE2 was knockdown using GIPZ lentiviral shRNA constructs (Horizon Discovery). Mature antisense sequences used for gene knockdown, are as follows:

shRNA Non-silencing control #RHS4346: ATCTCGCTTGGGCGAGAGTAAG

shRNA NSMCE2 #V2LHS_179143: TACATAATGGTTTAGTTGC

shRNA NSMCE2 #V3LHS_354626: TATTGTAGATTGAACAGCC

shRNA NSMCE2 #V3LHS_354628: ATTTTGAAAGTCTGCATCA

Lentiviral particles were generated by co-transfection of 293T cells with the packaging plasmids psPAX2 and pMD2G, and a mix of the three pGIPZ-shNSMCE2 specific constructs or the pGIPZ-shControl plasmid using TurboFect Transfection Reagent (Life Technologies). Lentiviral particles were harvested 48 hours after transfection in the culture media and used to infect target cells. After 24 hours incubation, transduced cells were selected with puromycin. Knockdown efficiency was quantified by qPCR.

## Statistical Analysis

All experiments were done in triplicates unless otherwise indicated. Results were plotted as mean values +/- standard deviation using GraphPad Prism7 and statistically analyzed using ANOVA followed by Dunnet’s or Bonferroni’s post-test, as appropriate. P-values less than 0.05 were considered statistically significant.

## Data availability

The dbGAP accession number for the PDX breast tumour H3K27ac ChIP-Seq data sets previously generated is phs001264.v1.p1 ^18^.

