## supplementary figures and tables for "NSMCE2, a Novel Super-Enhancer Regulated Gene, is Linked to Poor Prognosis and Therapy Resistance in Breast Cancer"

Supplementary Figure 1

A)

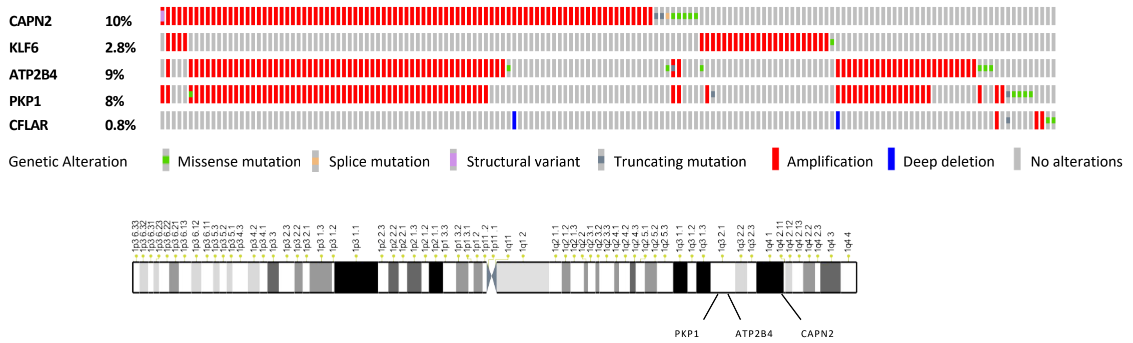


B)

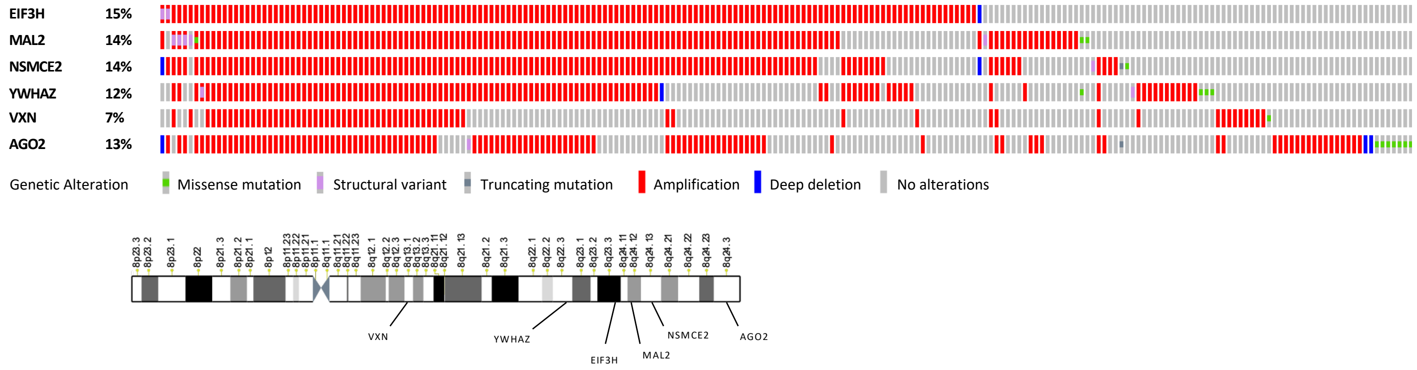


**Supplementary Figure 1**

Top figures are oncoprints showing genetic alterations identified in breast primary tumor samples (each bar represents one sample) for: A) genes grouped in Cluster 1 and B) genes grouped in Cluster 4 by our gene correlation matrix. Bottom figures are chromosome representation showing gene locations for: A) genes in Cluster 1 and B) genes in Cluster 4. Oncoprints were built on cBioPortal using TCGA breast cancer data. Red bars show DNA amplifications, blue bars show DNA deletions, and grey bars with green, orange, purple or dark grey marks show DNA mutations. Samples that do not have genetic alterations (gray bars) in any of the genes analyzed are not shown.

Supplementary Figure 2

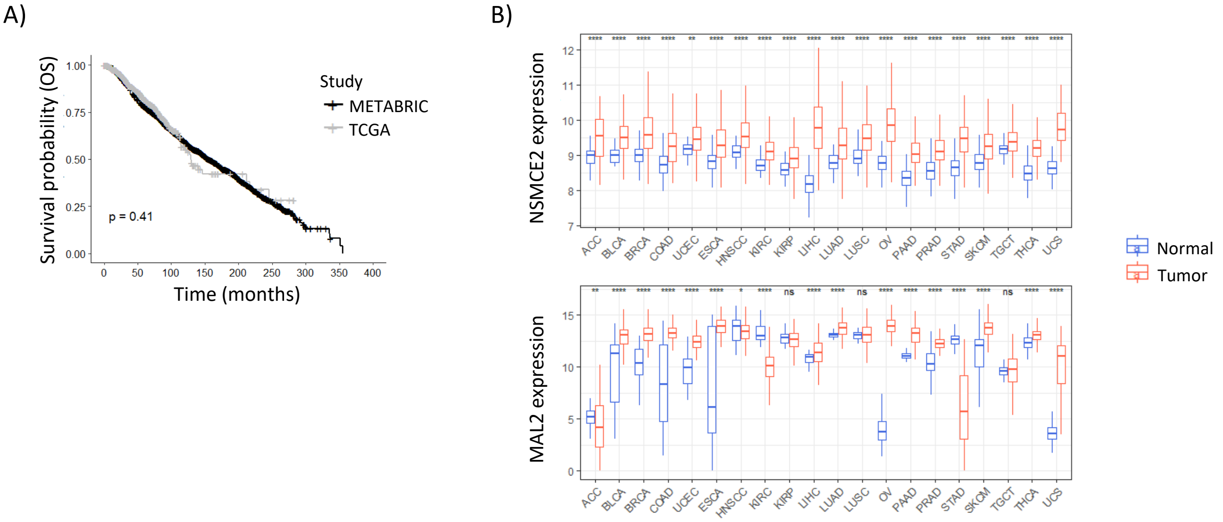


**Supplementary Figure 2**

A) Kaplan-Meier plots showing survival probability over time for breast cancer patients on TCGA (gray curve, n = 1097) and on METABRIC (black curve, n = 1904). Log-rank test was performed. B) Box plots showing NSMCE2 and MAL2 RNA-seq expression in normal versus tumor samples for pan-cancer analysis. Normal tissue includes data from GTEX and TCGA Solid Tissue Normal. Tumor sample data is from TCGA Pan-Cancer. Wilcoxon test was performed, ns = non-significant, *P < 0.05, **P<0.01, ***P<0.001, ****P<0.0001. OS = overall survival, ACC= Adrenocortical Cancer, BLCA=Bladder Urothelial Carcinoma, BRCA= Breast Invasive Carcinoma, COAD= Colon Adenocarcinoma, UCEC= Uterine Corpus Endometrioid Carcinoma, ESCA= Esophageal Carcinoma, HNSCC= Head and Neck Squamous Cell Carcinoma, KIRC= Kidney Clear Cell Carcinoma, KIRP= Kidney Papillary Cell Carcinoma , LIHC= Liver Hepatocellular Carcinoma, LUAD= Lung Adenocarcinoma, LUSC = Lung Squamous Cell Carcinoma, OV= Ovarian Serous Cystadenocarcinoma, PAAD= Pancreatic Ductal Adenocarcinoma, PRAD= Prostate Adenocarcinoma, STAD= Stomach Adenocarcinoma, SKCM= Skin Cutaneous Melanoma, TGCT= Testicular Germ Cell Tumor, THCA= Thyroid carcinoma, UCS= Uterine Carcinosarcoma.

Supplementary Figure 3

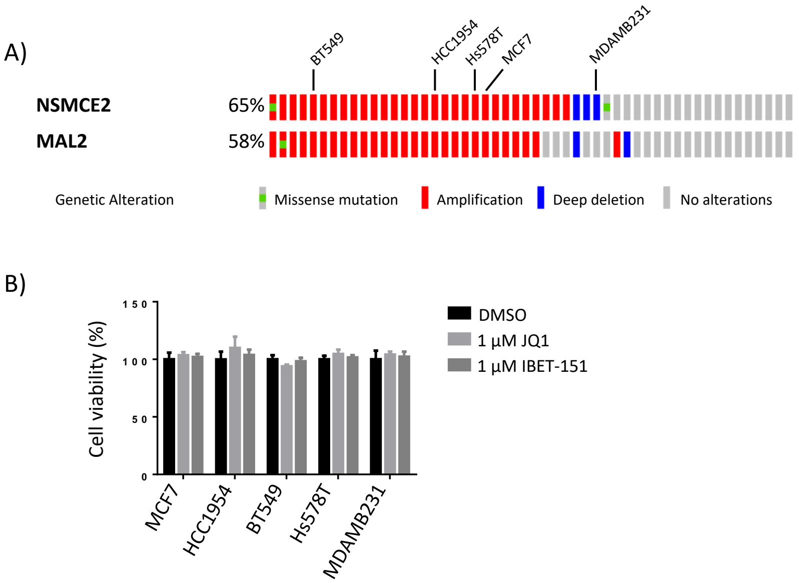


**Supplementary Figure 3**

A) Oncoprints showing breast cancer cell lines with genetic alterations for NSMCE2 and MAL2. Oncoprints were built on cBioPortal using Cancer Cell Line Encyclopedia dataset. Each column represents one cell line. Red samples show DNA amplifications, blue samples show DNA deletions, green dots show missense mutations. Samples that do not have genetic alterations are shown in gray. B) Effect of BET inhibition on cell viability of MCF7, HCC1954, BT-549, Hs578T and MDA-MB-231 measured by MTT assay. Cell lines were treated with vehicle (DMSO), 1 µM JQ1 or 1 µM IBET-151 for 24 hours.

Supplementary Figure 4

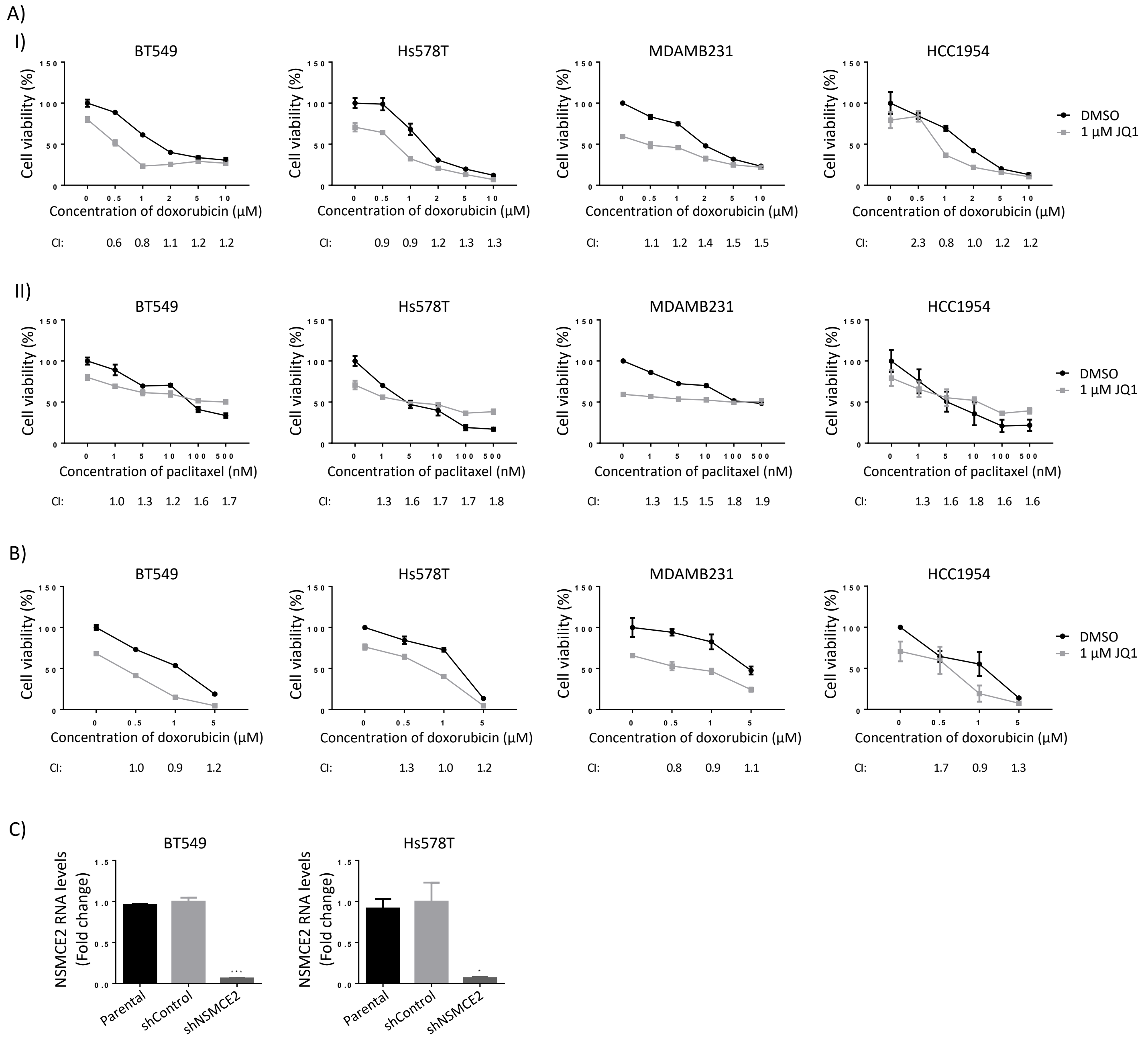


**Supplementary Figure 4**

A) Effect of combination of JQ1 and chemotherapeutic drugs on cell viability of breast cancer cell lines. BT-549, Hs578T, MDA-MB-231, and HCC1954 were preincubated with vehicle (DMSO) or 1 µM JQ1 for 3 hours, after which culture medium was replaced by fresh medium containing vehicle or 1 µM JQ1 and the indicated concentrations of (I) doxorubicin or (II) paclitaxel. After 48 hours, viable cells were quantified by the MTT assay. The Combination index (CI) was calculated for each combined treatment and is shown below each chemotherapeutic agent concentration. CI = 1 indicates additive effect, CI < 1 indicates synergistic effect, and CI > 1 indicates antagonistic effect. B) Effect of combination of JQ1 and doxorubicin on cell viability of breast cancer cell lines. BT-549, Hs578T, MDA-MB-231, and HCC1954 were preincubated with vehicle (DMSO) or 1 µM JQ1 for 24 hours, after which culture medium was replaced by fresh medium containing vehicle or 1 µM JQ1 and the indicated concentrations of doxorubicin. After 48 hours, viable cells were quantified by the MTT assay. The Combination index (CI) was calculated for each combination treatment and is shown below each doxorubicin concentration. CI = 1 indicates additive effect, CI < 1 indicates synergistic effect, and CI > 1 indicates antagonism. C) Analysis of NSMCE2 knockdown efficiency in breast cancer cell lines. BT549 and Hs578T cell lines were transduced with lentiviral particles for the expression of small hairpin RNAs against NSMCE2 (shNSMCE2) or control (shControl) and, after selection of cells that incorporated the transduced DNA with puromycin, NSMCE2 silencing efficiency was determined by qPCR. Parental cells were included in the qPCR to assess the effect of shControl on NSMCE2 expression. ANOVA followed by Dunnett's multiple comparison test was performed, *P<0.05, ***P<0.001.

Supplementary Table 1

| Official symbol | Official full name | Chromosome | Association with OS | P-value KM plot OS | Expression in GTEX | Expression in Normal TCGA | Expression in Tumor TCGA |
|---|---|---|---|---|---|---|---|
| SNORA14B | small nucleolar RNA, H/ACA box 14B | 1 | | NA | - | - | - |
| PKP1 | plakophilin 1 | 1 | | ns | 10.7 | **11.3** | 8.43 |
| MACF1 | microtubule actin crosslinking factor 1 | 1 | | ns | 13.3 | **13.9** | 13.3 |
| PHLDA3 | pleckstrin homology like domain family A member 3 | 1 | | ns | **11.7** | 11.0 | 10.5 |
| CAPN2 | calpain 2 | 1 | | ns | **13.6** | 13.2 | 13.1 |
| ATP2B4 | ATPase plasma membrane Ca2+ transporting 4 | 1 | | ns | 12.6 | **13.2** | 12.3 |
| MIR4435-2HG | MIR4435-2 host gene | 2 | | ns | 9.02 | 8.70 | **9.56** |
| CYTOR | cytoskeleton regulator RNA | 2 | | ns | - | - | - |
| CFLAR | CASP8 and FADD like apoptosis regulator | 2 | | ns | **13.3** | 12.7 | 11.6 |
| NDST1 | N-deacetylase and N-sulfotransferase 1 | 5 | | ns | 12.2 | **13.2** | 12.5 |
| LINC02532 | long intergenic non-protein coding RNA 2532 | 6 | | NA | - | - | - |
| NSMCE2 | NSE2 (MMS21) homolog, SMC5-SMC6 complex SUMO ligase | 8 | Bad prognosis | 0.003558 | 9.14 | 8.91 | **9.59** |
| VXN | vexin | 8 | | ns | 5.81 | **7.15** | 4.80 |
| TRPS1 | transcriptional repressor GATA binding 1 | 8 | | ns | 13.0 | 12.1 | **13.8** |
| EIF3H | eukaryotic translation initiation factor 3 subunit H | 8 | Bad prognosis | 0.01526 | 13.1 | 12.6 | **13.4** |
| MAL2 | mal, T cell differentiation protein 2 | 8 | Bad prognosis | 0.001970 | 11.9 | 9.75 | **13.1** |
| YWHAZ | tyrosine 3-monooxygenase/tryptophan 5-monooxygenase activation protein zeta | 8 | | ns | 14.6 | 13.8 | **15.2** |
| AGO2 | argonaute RISC catalytic component 2 | 8 | | ns | 11.2 | **11.3** | 10.9 |
| KLF6 | Kruppel like factor 6 | 10 | | ns | **13.9** | 13.8 | 12.4 |
| CUEDC1 | CUE domain containing 1 | 17 | | ns | 10.1 | 10.6 | **10.7** |
| LINC00673 | long intergenic non-protein coding RNA 673 | 17 | | NA | - | - | - |
| ANKRD12 | ankyrin repeat domain 12 | 18 | | ns | 11.3 | **11.5** | 10.9 |
| DLGAP1-AS1 | DLGAP1 antisense RNA 1 | 18 | Good prognosis | 0.004122 | 7.41 | 7.14 | **7.49** |
| BCAS4 | breast carcinoma amplified sequence 4 | 20 | | ns | 7.69 | 8.07 | **9.72** |
| ATP9A | ATPase phospholipid transporting 9A (putative) | 20 | | ns | 11.4 | **12.6** | 12.1 |
| SUMO1P1 | SUMO1 pseudogene 1 | 20 | | ns | - | - | - |

**Supplementary Table 1.**

**Gene expression values for SE-associated genes in different breast sample types and their correlation to survival probability.** Table showing official symbol, full name, chromosome location, association of high gene expression with OS observed in Kaplan Meier plots, p-values of Kaplan Meier plot, and gene expression levels in healthy tissue samples (GTEX), normal adjacent tissue (TCGA normal), and primary breast tumors (TCGA) of candidate SE-associated genes. Kaplan Meier plots were done using RNA-seq expression data on primary breast tumors on TCGA and analyzed by the Log-rank test. OS = overall survival, ns= non-significant, NA = non-available.

Supplementary Table 2

| Cancer Type | P-value NSMCE2 | P-value MAL2 |
|---|---|---|
| Breast Invasive Carcinoma | 0.0036 | 0.002 |
| Pancreatic Ductal Adenocarcinoma | 0.0447 | 0.0011 |
| Uveal Melanoma | 0.0058 | 0.0219 |
| Kidney Papillary Cell Carcinoma | 0.0013 | ns |
| Lung Adenocarcinoma | 0.0301 | ns |
| Sarcoma | 0.0378 | ns |
| Brain Lower Grade Glioma | 0.0207 | ns |
| Thymoma | ns | 0.0455 |
| Uterine Corpus Endometrial Carcinoma | ns | 0.0121 |
| Bladder Urothelial Carcinoma | ns | 0.0421 |
| Head and Neck Squamous Cell Carcinoma | ns | 0.033 |
| Cervical & Endocervical Cancer | ns | ns |
| Acute Myeloid Leukemia | ns | na |
| Adrenocortical Cancer | ns | ns |
| Colon Adenocarcinoma | ns | ns |
| Esophageal Carcinoma | ns | ns |
| Kidney Clear Cell Carcinoma | ns | ns |
| Liver Hepatocellular Carcinoma | ns | ns |
| Lung Squamous Cell Carcinoma | ns | ns |
| Ovarian Serous Cystadenocarcinoma | ns | ns |
| Prostate Adenocarcinoma | ns | ns |
| Rectum Adenocarcinoma | ns | ns |
| Skin Cutaneous Melanoma | ns | ns |
| Stomach Adenocarcinoma | ns | ns |
| Testicular Germ Cell Tumor | ns | ns |
| Thyroid Carcinoma | ns | ns |
| Uterine Carcinosarcoma | ns | ns |

In red: High gene expression associates with shorter survival
In blue: High gene expression associates with longer survival

**Supplementary Table 2**

**Association between high NSMCE2 or MAL2 expression levels and survival in different cancer types.** p-values correspond to Log-rank test for Kaplan-Meier plots for overall survival probability over time based on NMSCE2 and MAL2 RNA-seq expression on TCGA Pan-Cancer data. Red values denote high gene expression associates with shorter survival, blue values denote high gene expression associates with longer survival. ns = non-significant, na = non-available.

Supplementary Table 3

| Official symbol | pCR to Chemotherapy in BC patients (n=507) | | | | pCR to Chemotherapy in Grade III BC patients (n=194) | | | |
|---|---|---|---|---|---|---|---|---|
| | AUC | ROC p-value | Mann-Whitney test p-value | Higher expression in patient group | AUC | ROC p-value | Mann-Whitney test p-value | Higher expression in patient group |
| MIR4435-2HG | 0.524 | 0.22 | 0.42 | Responder | 0.528 | 0.29 | 0.55 | Responder |
| SNORA14B | NA | NA | NA | NA | NA | NA | NA | NA |
| ANKRD12 | 0.515 | 0.17 | 0.34 | Responder | 0.528 | 0.13 | 0.25 | Responder |
| CYTOR | NA | NA | NA | NA | NA | NA | NA | NA |
| PKP1 | 0.567 | 5.60E-06 | 1.10E-05 | Responder | 0.501 | 49 | 0.98 | Non-responder |
| NSMCE2 | 0.617 | 0.000032 | 0.00011 | Non-responder | 0.655 | 0.00017 | 0.00099 | Non-responder |
| MACF1 | 0.561 | 3.90E-05 | 7.00E-05 | Responder | 0.513 | 0.29 | 0.58 | No change |
| LINC02532 | NA | NA | NA | NA | NA | NA | NA | NA |
| VXN | 0.547 | 4.50E-02 | 0.12 | Responder | 0.53 | 0.25 | 0.52 | Responder |
| LINC00673 | NA | NA | NA | NA | NA | NA | NA | NA |
| BCAS4 | 0.502 | 0.48 | 0.95 | No change | 0.506 | 0.44 | 0.89 | Responder |
| PHLDA3 | 0.594 | 4.80E-11 | 6.50E-10 | Non-responder | 0.509 | 0.35 | 0.69 | Responder |
| KLF6 | 0.529 | 0.18 | 0.33 | Non-responder | 0.568 | 0.072 | 0.15 | Non-responder |
| CAPN2 | 0.548 | 7.10E-04 | 0.0016 | Responder | 0.532 | 0.093 | 0.19 | Non-responder |
| TRPS1 | 0.513 | 0.21 | 0.4 | Responder | 0.573 | 0.00088 | 0.0024 | Non-responder |
| CFLAR | 0.559 | 3.00E-02 | 0.05 | Responder | 0.501 | 0.49 | 0.98 | Non-responder |
| EIF3H | 0.511 | 0.24 | 0.49 | No change | 0.545 | 0.027 | 0.061 | Non-responder |
| ATP9A | 0.516 | 1.50E-01 | 0.3 | Non-responder | 0.544 | 0.034 | 0.067 | Non-responder |
| CUEDC1 | 0.523 | 0.23 | 0.44 | Responder | 0.536 | 0.25 | 0.45 | Responder |
| MAL2 | 0.557 | 0.003 | 0.06 | Non-responder | 0.554 | 0.13 | 0.25 | Non-responder |
| NDST1 | 0.533 | 0.013 | 0.029 | Responder | 0.553 | 0.015 | 0.028 | Responder |
| DLGAP1-AS1 | 0.542 | 0.08 | 0.16 | Responder | 0.63 | 0.0017 | 0.0055 | Responder |
| ATP2B4 | 0.503 | 0.41 | 0.82 | No change | 0.54 | 0.045 | 0.094 | Non-responder |
| YWHAZ | 0.609 | 3.50E-14 | 7.20E-13 | Responder | 0.539 | 0.055 | 0.11 | Responder |
| AGO2 | 0.513 | 0.33 | 0.66 | No change | 0.57 | 0.052 | 0.13 | Non-responder |
| SUMO1P1 | NA | NA | NA | NA | NA | NA | NA | NA |

**Supplementary Table 3**

**Association between RNA expression of SE-associated genes and response to chemotherapy in breast cancer patients.** Values for AUC, ROC p-value and Mann-Whitney test p-value are shown for each gene, along with the patient group with higher gene expression in breast cancer patients (n = 507) and in Grade III breast cancer patients (n = 194). The analysis was performed using ROC Plotter. pCR = pathological complete response, NA = non-available.
